# PaReBrick: PArallel REarrangements and BReakpoints identification toolkit

**DOI:** 10.1101/2021.05.18.444676

**Authors:** Alexey Zabelkin, Yulia Yakovleva, Olga Bochkareva, Nikita Alexeev

## Abstract

**Motivation:** High plasticity of bacterial genomes is provided by numerous mechanisms including horizontal gene transfer and recombination via numerous flanking repeats. Genome rearrangements such as inversions, deletions, insertions, and duplications may independently occur in different strains, providing parallel adaptation. Specifically, such rearrangements might be responsible for multi-virulence, antibiotic resistance, and antigenic variation. However, identification of such events requires laborious manual inspection and verification of phyletic pattern consistency.

**Results:** Here we define the term “parallel rearrangements” as events that occur independently in phylogenetically distant bacterial strains and present a formalization of the problem of parallel rearrangements calling. We implement an algorithmic solution for the identification of parallel rearrangements in bacterial population, as a tool PaReBrick. The tool takes synteny blocks and a phylogenetic tree as input and outputs rearrangement events. The tool tests each rearrangement for consistency with a tree, and sorts the events by their parallelism score and provides diagrams of the neighbors for each block of interest, allowing the detection of horizontally transferred blocks or their extra copies and the inversions in which copied blocks are involved. We proved PaReBrick’s efficiency and accuracy and showed its potential to detect genome rearrangements responsible for pathogenicity and adaptation in bacterial genomes.

**Availability:** PaReBrick is written in Python and is available on *[GitHub](https://github.com/ctlab/parallel-rearrangements)*

## 1 Introduction

Bacterial chromosomes contain numerous repeats ranging from short motifs to gene paralogs, which provide substrates for recombination affecting genome composition and gene order [4]. Such rearrangements make a significant contribution to the evolution of prokaryotic species since they can lead to the formation of new genes, disruption of existing genes, and modulation of expression level [14]. Large-scale deletions and inversions affect different levels of chromosome organization, and are mostly deleterious [16, 3].

Nevertheless, beneficial rearrangements are also known, such as the ones providing acquisition of new functions, phenotype switching, or fast genome reduction [2]. Such rearrangements may occur independently in different strains providing parallel adaptation to new environments [17] or phenotypic diversity into clonal populations [20]. The latter is shaped by phase variation — the mechanism of reversible alternation between genetic states. A strategy that may allow fast and cheap prediction of putative mechanisms of variation is the computational analysis of genomes focusing on genomic repeats and screening for reversible genome rearrangements. For instance, in the human pathogen *Streptococcus pneumoniae*, reversible inversions affecting surface antigens, encoded by *phtB* and *phtD* genes, were revealed computationally and confirmed experimentally [18, 19].

However, identification of rearrangements responsible for parallel evolution or phenotype switching requires laborious manual inspection of the composition and order of synteny blocks across the species’ phylogenetic tree and verification of the consistency of evolutionary events with tree topology [1, 18, 17]. Here we introduce PaReBrick, a method to identify and visualise parallel rearrangements in bacterial populations. We proved PaReBrick’s efficiency and accuracy and showed its potential to detect genome rearrangements responsible for pathogenicity and adaptation in bacterial genomes.

## 2 Methods

### 2.1 Data preprocessing

#### Phylogenetic tree

Nowadays there are a lot of implemented approaches to construct phylogenetic trees which are usually based on concatenated alignment of homologous genes. In our study we use the PanACoTA pipeline [15] which includes all steps for phylogenetic tree construction including genome annotation and orthologs detection. Thus, intermediate results can be further used for biological annotation and interpretation.

#### Synteny blocks

In our study, we understand by *synteny blocks* a decomposition of genomes into non-overlapping highly conserved segments. Synteny blocks can be defined on different scales depending on the field of study, and the scale is usually controlled by the threshold of minimal block length. Synteny blocks are often constructed based on *seeds* (also called *anchors*), with most methods using the “seed-and-extend” approach. Homologous genes, locally-collinear blocks or any multiple whole-genome alignment results can be used as seeds. In our study we use the efficient multiple whole-genome alignment tool SibeliaZ [13] to obtain locally collinear blocks and its submodule `maf2synteny` [11] to construct synteny blocks.

### 2.2 What is parallel?

We say that a rearrangement is *consistent* with a tree if we may associate it with a particular branch on a tree, otherwise we call a rearrangement *parallel* (Fig. 1e). More formally, a *character* is consistent with a tree if any two strains sharing a character state have a common ancestor with the same character state. We test each character for consistency with a tree by solving the maximum parsimony problem [7, 6].

**Figure 1:**
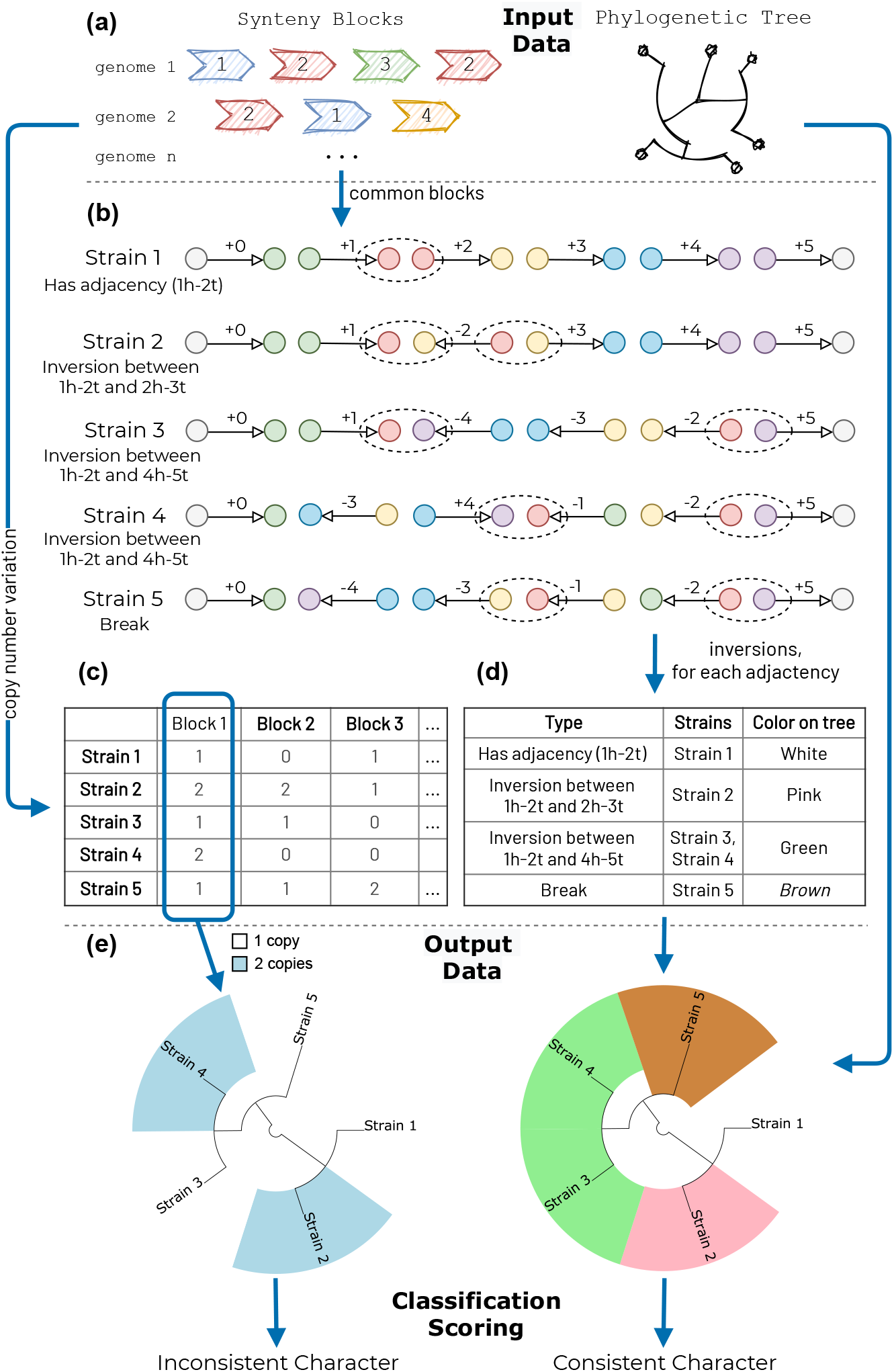
The PaReBrick pipeline: **(a)** the tool takes a representation of synteny blocks in the strains and their phylogenetic tree; analyzes **(b)** balanced and **(c)** unbalanced rearrangements; and **(d)** assigns a character state to each strain based on this analysis. For each character the tool tests if it is consistent with the tree **(e)**; if not, it computes the parallelism score and reports all the inconsistent events.

Not all parallel rearrangements are equally significant, so we introduce a parallelism score (see Section 3.2). For example, in Fig. S1a the rearrangement seems to happen independently twice, but the corresponding nodes in a tree are close to each other, so the inconsistency may be explained by artifacts in the input data. In contrast, in Fig. S1d the rearrangement occurred independently in several distant nodes, so there is more evidence that this is an actual parallel event.

We analyze two classes of genome rearrangements: *balanced*, which change the order of synteny blocks, but do not delete or duplicate them, and *unbalanced*, which affect block copy numbers, such as indels and duplications. We associate a character with each rearrangement and assign each strain a character state based on this.

## 3 Results

The proposed tool takes synteny blocks in multiple genomes and a phylogenetic tree as an input, and it outputs the list of rearrangement events sorted by their parallelism score (Fig. 1).

### 3.1 Characters assignment

#### Characters for unbalanced rearrangements

To analyse indels and duplications, we consider all blocks which were present in different copy numbers in the strains. We associate a character with each block *B* and assign to each strain a character state equal to the number of copies of *B* presented in this strain (Fig. 1c).

Genome rearrangements may occur on different evolutionary scales and overlap. For example, a large insertion may be followed by a short deletion, resulting in similar but not identical occurrence patterns of the involved blocks. PaReBrick automatically clusters the blocks according to the similarity of their occurrence patterns (see Section 3.3).

#### Characters for balanced rearrangements

In order to analyze balanced rearrangements, we concentrate on common single-copy block content, that is we consider only the those blocks present in each strain exactly once.

We represent each strains’ genome as a circular sequence of synteny blocks. Each block (say, block 1) is represented by its tail (1*t*) and head (1*h*). Consecutive blocks (say, blocks 1 and 2) are linked by an *adjacency* (1*h -* 2*t*) (Fig. 1b).

We associate a character with each adjacency and assign the character state to each strain according to the classification below:

1. adjacency is present;
2. adjacency is absent, and the breakage can be explained by a single inversion;
3. adjacency is absent, and the breakage is the result of multiple rearrangements

### 3.2 Parallelism score

In this section we interpret character states as colors for better readability.

To range the characters, we start with the reconstruction of the character colors in the internal nodes of the phylogenetic tree. For this step, we use a modification of the algorithm introduced by [6] based on a dynamic programming approach.

After that, for each color *c* we find the set of all the vertices *V*_*c*_ on a tree where it changed: *V*_*c*_ = {*v* |color(*v*) = *c* & color(ancestor(*v*)) ≠ *c*}.

Let’s say that some *color is inconsistent* with a tree if it has at least two independent appearances. See Supplements Appendix A for examples. Formally, *c* — inconsistent if |*V*_*c*_| > 1.

We note that the inconsistent character is a character with at least one inconsistent color.

Color inconsistency is calculated as the sum of the distances between all pairs of independent appearances of some color:

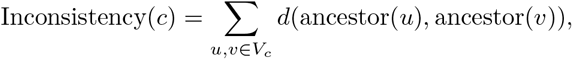

where *d*(*u, v*) — is distance between nodes on a phylogenetic tree (see example on Fig. 3).

**Figure 2:**
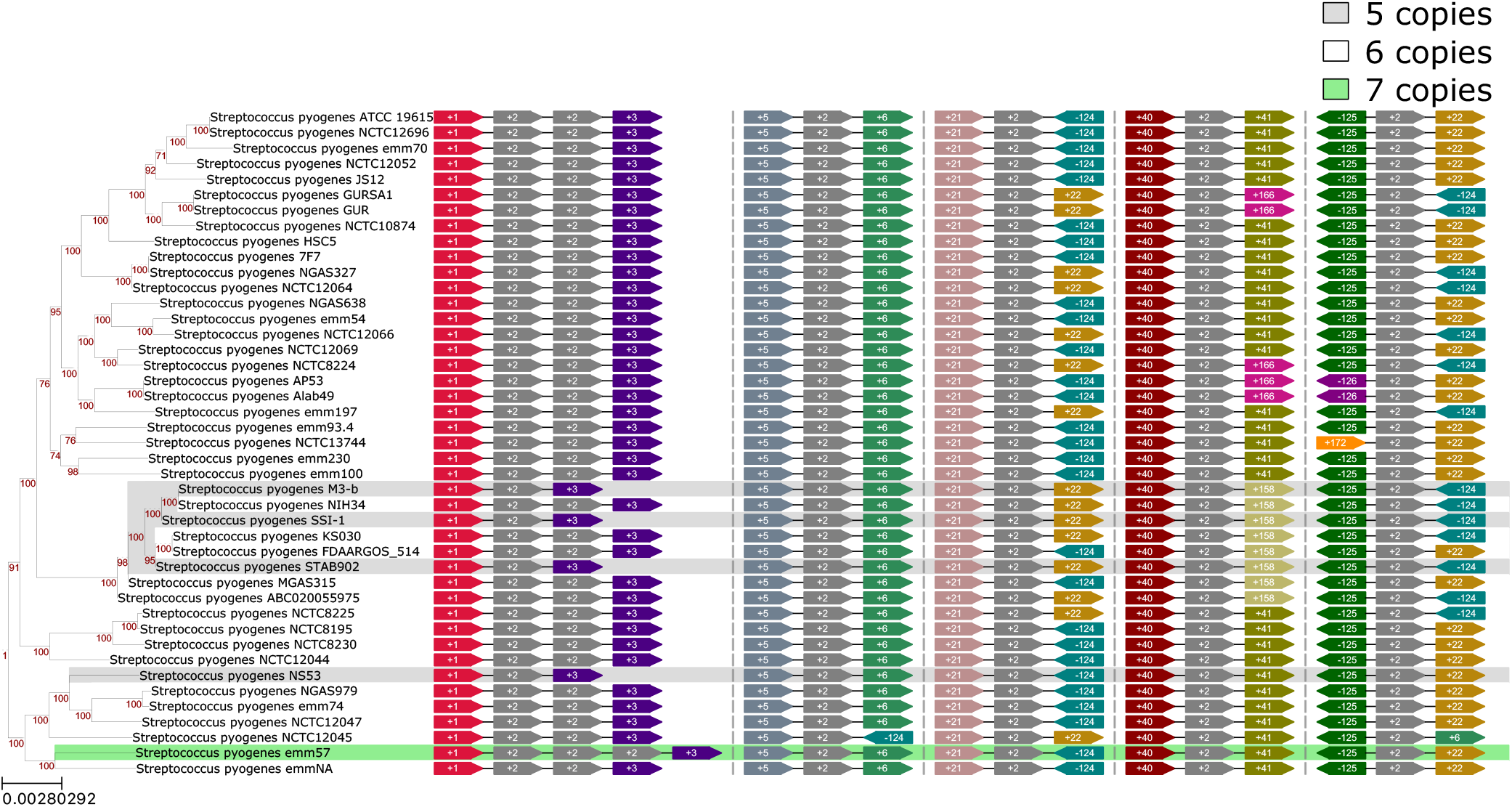
The genomic context of **block** #2, comprising the rRNA gene operon. For better visibility the subtree is used. For each strain and each copy of the block, the upstream and the downstream neighbouring block are shown. It revealed an variation of number of tandem copies in the first loci and a reversible inversion between copies in the third and the fifth loci.

**Figure 3:**
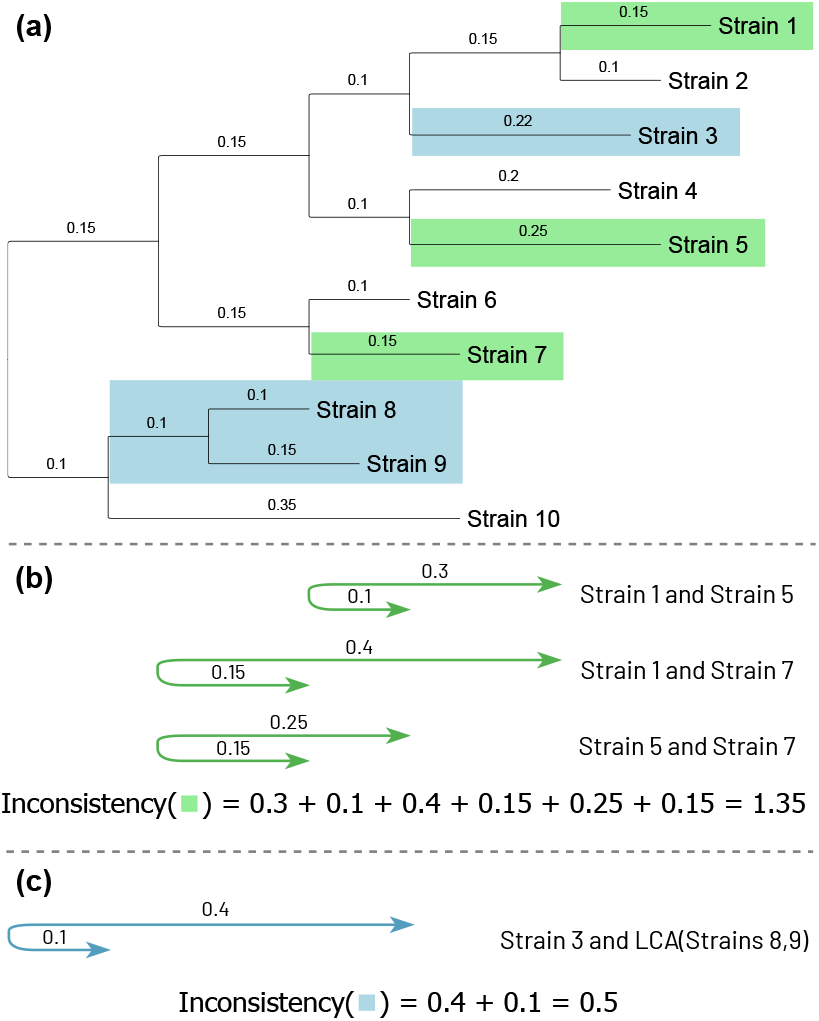
Calculation of inconsistency of colors. **(a)** The phylogenetic tree is coloured to reflect states of the feature **(b**,**c)** For each color, inconsistency is calculated as a sum of the distances between all pairs of independent appearances of that color.

We use the following metrics to estimate the parallelism score of a character:

1. *Parallel rearrangement score*: ∑_*c*_ Inconsistency(*c*), sum of inconsistencies for all colors;
2. *Number of inconsistent colors* : #{*c* : |*V*_*c*_| > 1};
3. *Total number of parallel events* : 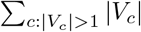, number of appearances for all inconsistent colors.

We also introduce the *break score* to rank the rearrangements resulted from multiple breakages, see Supplements Appendix A.2 for details.

### 3.3 Clustering of unbalanced rearrangements by phyletic patterns

To cluster the blocks, we take into account two features:

1. *Similarity of tree patterns*. With each block *b* we associate a vector *v*(*b*) of length *n*, where *n* is the total number of strains, in the following way: *v*_*k*_(*b*) is the copy number of the block *b* in the *k*-th strain. As a measure of dissimilarity between block patterns for blocks *b*_1_ and *b*_2_ we use the Manhattan

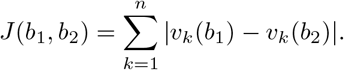
2. *Distance between blocks*. For each pair of blocks we can calculate the minimal distance between those blocks for every strain. As a measure of distance *B*(*b*_1_, *b*_2_) we take the first quartile of this distribution. If some blocks *b*_1_ and *b*_2_ don’t appear together in any strain, we assign the longest genome length to *B*(*b*_1_, *b*_2_).

Further, the matrices *J* and *B* are normalized so that all their values are in the interval [0, 1]. Then we apply hierarchical clustering to the blocks based on a distance matrix *D* = *j* · *J* + *b* · *B* (by default *j* = 0.8, *b* = 0.2).

### 3.4 Neighborhood visualization

As various molecular mechanisms might be responsible for variations in block content, we provide a diagram of the neighbors for each block in each genome where it is present (See Figure 2). For easier comparison of their context in different genomes, all blocks’ copies are rotated on the same side and grouped into columns based on similarity of their neighbours. If tandem copies of a block are present, all copies are visualized. This visualization aims for the best readability of blocks’ context data and therefore does not reflect the order of the loci in genomes nor blocks’ length. Meanwhile, it allows to detect horizontally transferred blocks and to distinguish between copies of a block. For multi-copied blocks, this visualization reveals inversions in which these repeats are involved.

## 4 Application

To prove the method efficiency and accuracy, we applied the PaReBrick tool to complete genomes of *Streptococcus pneumoniae*. The PaReBrick tool detected the inversion between *PhtB* and *PhtD* genes as an event with the highest parallelism score (Supplementary Fig.S3) This inversion was previously shown to be respon for antigenic variation affecting surface antigens, encoded by *PhtB* and *PhtD* genes [18, 19]. PaReBrick detected nine strains across the phylogenetic tree that have this fragment inverted, including five strains that were identified in [18]. Moreover, PaReBrick detected additional rearrangement events affecting these adjacencies in five strains.

Then we used the PaReBrick tool for investigation of parallel rearrangements in 219 *Streptococcus pyogenes* genomes (see Appendix B in Supplements). The complete assemblies of *S. pyogenes* were downloaded from the NCBI RefSeq database with the PanACoTA tool, all plasmids were excluded. The whole project including the input and output of the tool is available at the *[GitHub](https://github.com/ctlab/parallel-rearrangements)* repository.

We analysed 217 chromosomal synteny blocks, 125 of them present exactly once in each strain. Among six adjacencies involved in parallel rearrangements (Table 1) the highest parallelism score was assigned to a 1.4 Mb inversion (Fig. 4). In all strains, these adjacencies contain the multi-copied block with a *rRNA* operon, indicating its involvement in the recombination. The mean length of adjacencies (7 kb) is consistent with the operon length, which also validates the observation. These data closely resemble other previously described inversion events that have been shown to affect the underlying phenotype, including insertions of rRNA genes [10]. Thus, these data indicate that it is feasible to computationally detect parallel genomic events in closely related strains, possibly associated with phase variation in bacterial populations.

**Table 1:** Summary table for *balanced rearrangement* in *Streptococcus pyogenes*, only characters with *parallel rearrangement score* more than zero are shown. The first two lines in the table correspond to the inversion of the chromosome segment containing blocks #22…#124 in 52 strains. As the adjacencies 21h—22t and 124ht—125t were broken by the same rearrangement, they have the same scores.

| **Id** | **Adjacency** | **Mean**<br>**Break**<br>**Length**<br>**(nucleotide)** | **Parallel**<br>**Rearr.**<br>**Score** | **Number**<br>**of**<br>**Inconsistent**<br>**Colors** | **Number**<br>**of**<br>**Parallel**<br>**Events** | **Parallel**<br>**Break**<br>**Score** | **Number**<br>**of**<br>**Parallel**<br>**Breaks** |
| --- | --- | --- | --- | --- | --- | --- | --- |
| 1 | 124h—21h | 7894 | 6.57 | 3 | 51 | 16.27 | 53 |
| 2 | 125t—22t | 16751 | 6.37 | 2 | 49 | 15.35 | 52 |
| 3 | 101h—103t | 30744 | 0.013 | 1 | 2 | 0.13 | 5 |
| 4 | 51h—52t | 20481 | 0.013 | 1 | 2 | 0.086 | 4 |
| 5 | 135h—136t | 1665 | 0.0002 | 2 | 6 | 0.078 | 7 |
| 6 | 16h—17t | 3363 | 0.0002 | 2 | 6 | 0.0004 | 6 |
| ... |  |  |  |  |  |  |  |

**Figure 4:**
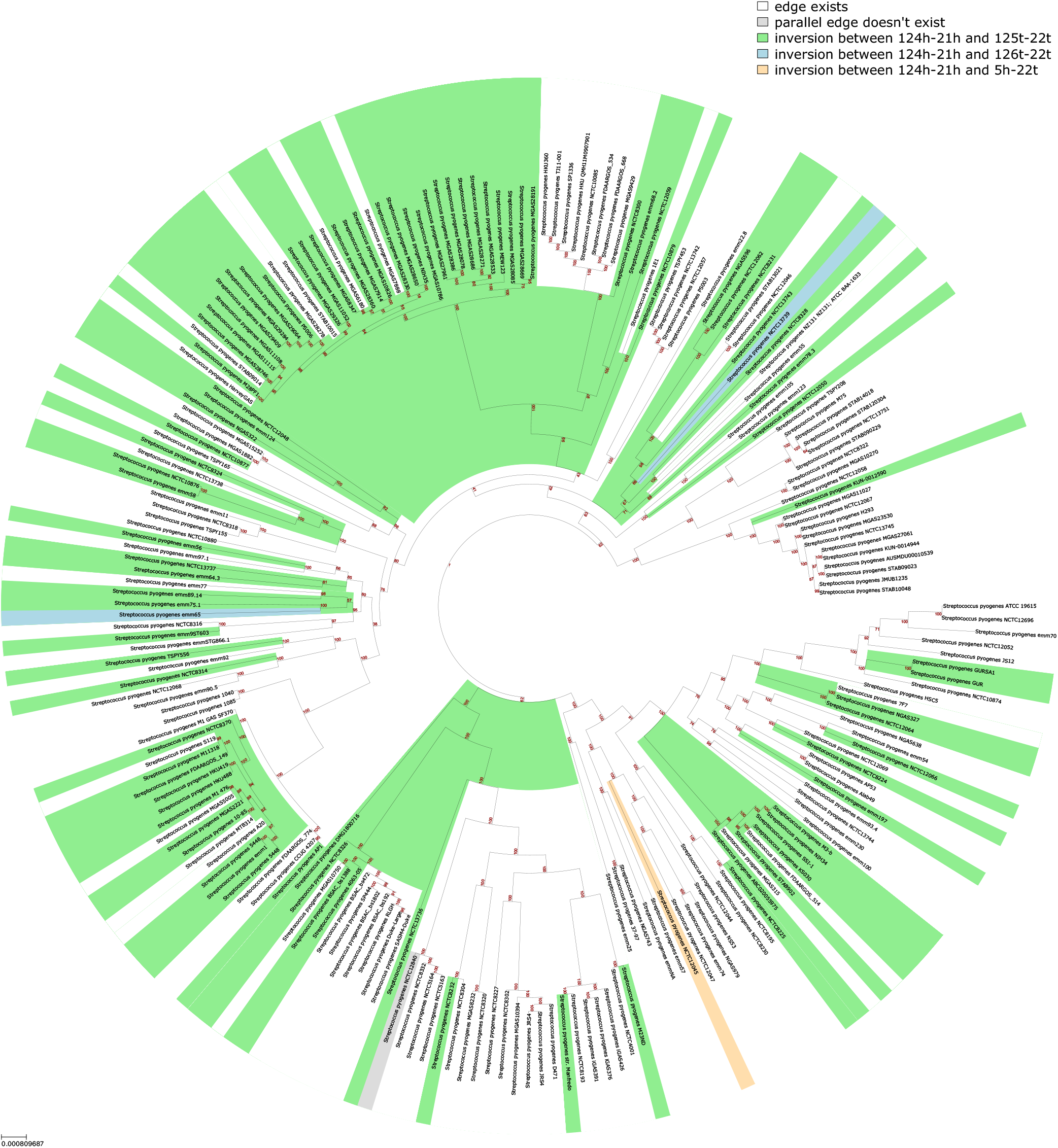
The tree is coloured to reflect states of the adjacency of the 124h and 21h syntenic blocks. White represents the ancestral state of the adjacency. In some strains the 1.4 Mb fragment is inverted between two copies of rRNA gene operons (green). In other strains the exact borders of this inversion are affected by additional events (orange and blue).

For 65 out of 217 blocks, we revealed parallel insertions, deletions, and multiplications (Table S1); the highest scores were assigned to phage insertions (Fig. S4). Visualization of genome context for these blocks revealed their independent acquisition by different *S. pyogenes* lineages. In some strains, these insertions occurred two or three times in different loci (Fig. S5). The block containing the *rRNA* operon also has high parallelism score. Indeed, while *S. pyogenes* genomes contain six copies of the block in five different loci, a few strains lost one of the tandem copies or gained up to four copies of the operon. (Figs. 2,S6,S7).

Visualization of blocks’ locations within this strain revealed that the reversible inversions occurred between a pair of repeats placed symmetrically across the origin (Fig. 5).

**Figure 5:**
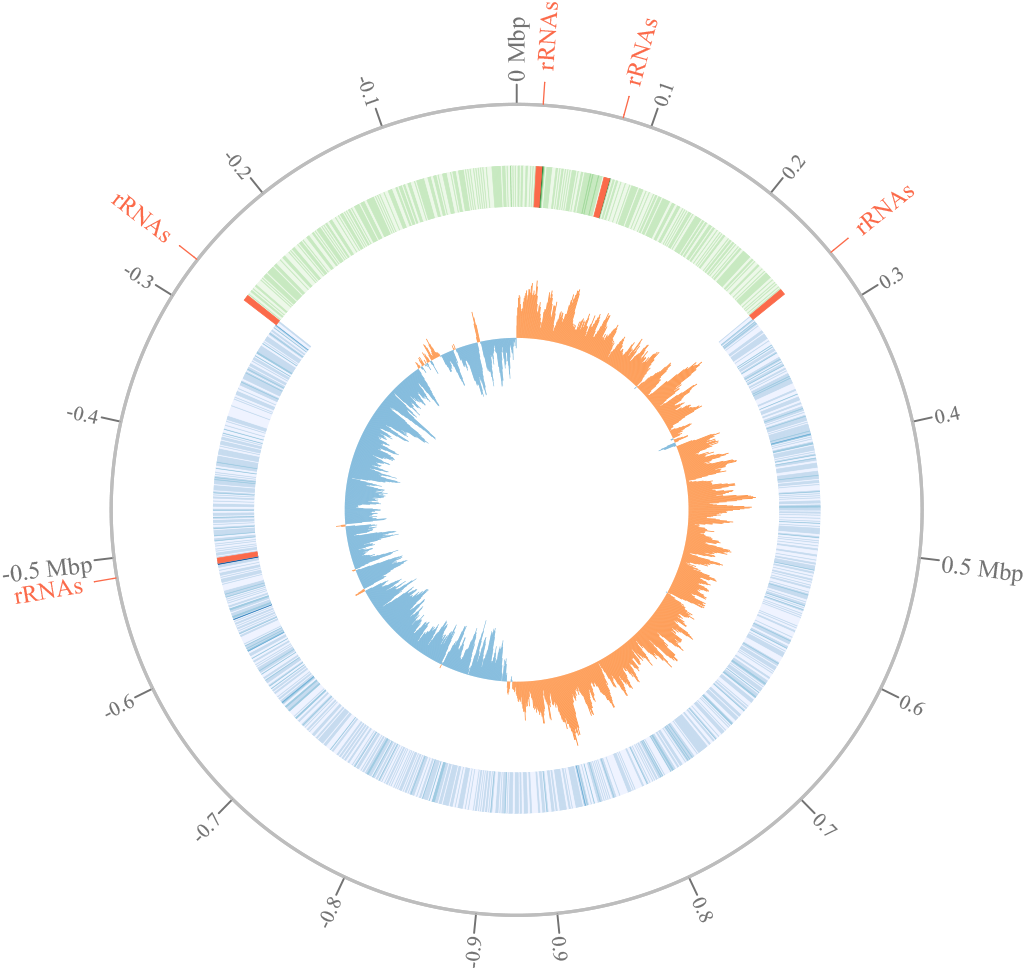
Relative positions of five loci containing rRNA gene operons in chromosomes of *Streptococcus pyogenes*. The reversible inversion typically occurs between a pair of symmetric repeats: for the presented *Streptococcus pyogenes* strain BSAC bs472 the distances from the origin to the repeats are 258,996 and 265,532 (based on rRNA block coordinates); and for 50% of all analyzed strains the corresponding repeats are located even more symmetrically. The inner blue-orange circle shows GC-skew, 0 Mbp corresponds to origin of replication (ori). The second circle represents the annotated genes, the green fraction of the chromosome is involved in reversible inversion. The borders of the inversion are formed by inverted copies of the rRNA gene operon located at the same distance from the ori.

## 5 Discussion

Our pipeline is based on a sequence of polynomial algorithms, and its complexity depends on the number of strains *s* and the number of blocks *b*. The step of constructing characters for unbalanced rearrangements has complexity *O*(*s*) for each block. Testing a character for consistency with a tree has complexity *O*(*s*) as well. The overall complexity of the clustering step is *O*(*b*^2^(*b*+*s*)) for building distance matrices and for hierarchical clustering itself. The step of constructing characters for balanced rearrangements takes *O*(*s*) for each adjacency. The total number of adjacencies is in between the number of blocks *b* (if all strains have same block order) and its square *b*^2^, but does not exceed *bs*.

In practice this means that for the *Streptococcus pyogenes* dataset with 219 strains the whole process takes 279 seconds on a laptop with Intel 4870HQ CPU, and the trees’ rendering consumes the majority of the computational time, 213 of 279 seconds.

Modern sequencing technology produces massive amounts of genomic data, providing exceptional opportunities to investigate whole-genome organization and interactions of different components [5, 12]. Several strategies are widely used for assembly validation such as long read (re)sequencing and/or PCR contiguity verification. Nevertheless, some of the detected rearrangements may been caused by inaccuracies in gap closure procedures. In turn, if the reference genome was used for gap resolving, some strain-specific genome rearrangements might be missed. Thus, for particular computational observations, further experimental validation may be required.

Genomic repeats of different nature may play the role of substrate for recombination. Recent studies have sparked a renewed interest in large-scale phase variation, as it may affect complex bacterial phenotypes and modulate expression of a set of genes[20]. Pathogenic bacterial species use this strategy for ensuring survival [9]. Phase variation might be responsible for chronic infections, providing multi-virulence, antibiotic resistance, and antigenic variation [19, 8, 10]. While particular cases are described, reversible large-scale inversions have been not investigated systematically. The PaReBrick tool allows for computational identification of phase variation. Systematization and verification of the observed cases is the key for understanding new molecular mechanisms in pathogens.

## 6 Conclusion

We proved PaReBrick’s efficiency and accuracy and showed its potential to detect genome rearrangements responsible for pathogenicity and adaptation in bacterial genomes. The PaReBrick tool has great potential to allow researchers to address wider research questions in evolutionary, molecular and medical fields. The approach might be used for the study of rapid emergence of new bacterial phenotypes, understanding the molecular basis of antibiotic resistance mechanisms and formation of small colony variants, and the study of the selective forces in genomic evolution underlying complex phenotypes. The application of this approach and the concomitant understanding of connections between detected genome rearrangements and medically-relevant phenotypes may contribute to the efficient development of drugs and vaccines.

## Acknowledgements

We thank the 2020 student class of the Bioinformatics Institute, who used the first versions of the tool and provided many valuable suggestions to improve usability. We also thank Louisa Gonzalez Somermeyer for manuscript proofreading.

## Funding

The work of AZ and NA were supported by National Center for Cognitive Research of ITMO University. The work of OB was supported by the European Union’s Horizon 2020 research and innovation programme under the Marie Sklodowska-Curie [754411].

## Notes

https://github.com/ctlab/parallel-rearrangements

### Competing Interest Statement

The authors have declared no competing interest.

## References

[1] Olga O. Bochkareva, Elena V. Moroz, Iakov I. Davydov, and Mikhail S. Gelfand. Genome rearrangements and selection in multi-chromosome bacteria Burkholderia spp. BMC Genomics, 19:965, 2018.

[2] G Brandis and D. Hughes. The snap hypothesis: Chromosomal rearrangements could emerge from positive selection during niche adaptation. PLoS Genet., 16(3):e1008615, 2020.

[3] AE Darling, I Miklós, and MA Ragan. Dynamics of genome rearrangement in bacterial populations. PLoS Genet, 4(7):e1000128, 2008.

[4] Elise Darmon and David R. F. Leach. Bacterial genome instability. Microbiology and Molecular Biology Reviews, 78(1):1–39, 2014.

[5] Adam C. English, Stephen Richards, Yi Han, Min Wang, Vanesa Vee, Jiaxin Qu, Xiang Qin, Donna M. Muzny, Jeffrey G. Reid, Kim C. Worley, and Richard A. Gibbs. Mind the gap: Upgrading genomes with pacific biosciences rs long-read sequencing technology. PLOS ONE, 7(11):1–12, 11 2012.

[6] Péter L. Erdős and László A. Székely. On weighted multiway cuts in trees. Mathematical Programming, 65(1-3):93–105, February 1994.

[7] Walter M. Fitch. Toward defining the course of evolution: Minimum change for a specific tree topology. Systematic Zoology, 20(4):406, December 1971.

[8] Romain Guérillot, Xenia Kostoulias, Liam Donovan, Lucy Li, Glen P. Carter, Abderrahman Hachani, Koen Vandelannoote, Stefano Giulieri, Ian R. Monk, Mayu Kunimoto, Lora Starrs, Gaétan Burgio, Torsten Seemann, Anton Y. Peleg, Timothy P. Stinear, and Benjamin P. Howden. Unstable chromosome rearrangements in staphylococcus aureus cause phenotype switching associated with persistent infections. Proceedings of the National Academy of Sciences, 116(40):20135–20140, September 2019.

[9] Xueting Huang, Juanjuan Wang, Jing Li, Yanni Liu, Xue Liu, Zeyao Li, Kurni Kurniyati, Yijie Deng, Guilin Wang, Joseph D Ralph, Megan De Ste Croix, Sara Escobar-Gonzalez, Richard J Roberts, Jan-Willem Veening, Xun Lan, Marco R Oggioni, Chunhao Li, and Jing-Ren Zhang. Prevalence of phase variable epigenetic invertons among host-associated bacteria. Nucleic Acids Research, 48(20):11468–11485, 10 2020.

[10] Sharon Irvine, Boyke Bunk, Hannah K. Bayes, Cathrin Spröer, James P. R. Connolly, Anne Six, Thomas J. Evans, Andrew J. Roe, Jörg Overmann, and Daniel Walker. Genomic and transcriptomic characterization of pseudomonas aeruginosa small colony variants derived from a chronic infection model. Microbial Genomics, 5(4), April 2019.

[11] M. Kolmogorov, B. Raney, B. Paten, and S. Pham. Ragout–a reference-assisted assembly tool for bacterial genomes. Bioinformatics, 30(12):i302–i309, jun 2014.

[12] M. A. Madoui, S. Engelen, C. Cruaud, C. Belser, L. Bertrand, A. Alberti, A. Lemainque, P. Wincker, and J. M. Aury. Genome assembly using nanopore-guided long and error-free dna reads. BMC Genomics, 16(1):327, 2015.

[13] Ilia Minkin and Paul Medvedev. Scalable multiple whole-genome alignment and locally collinear block construction with SibeliaZ. Nature Communications, 11(1), December 2020.

[14] Seema Patel. Drivers of bacterial genomes plasticity and roles they play in pathogen virulence, persistence and drug resistance. Infection, Genetics and Evolution, 45:151–164, November 2016.

[15] Amandine Perrin and Eduardo P C Rocha. PanACoTA: a modular tool for massive microbial comparative genomics. NAR Genomics and Bioinformatics, 3(1), 01 2021.

[16] Jelena Repar and Tobias Warnecke. Non-random inversion landscapes in prokaryotic genomes are shaped by heterogeneous selection pressures. Molecular Biology and Evolution, 34(8):1902–1911, April 2017.

[17] Zaira Seferbekova, Alexey Zabelkin, Yulia Yakovleva, Robert Afasizhev, Natalia Dranenko, Nikita Alexeev, Mikhail Gelfand, and Olga Bochkareva. High rates of genome rearrangements and pathogenicity of Shigella spp. Frontiers in Microbiology, 2021.

[18] Pavel V. Shelyakin, Olga O. Bochkareva, Anna A. Karan, and Mikhail S. Gelfand. Micro-evolution of three streptococcus species: selection, antigenic variation, and horizontal gene inflow. BMC Evolutionary Biology, 19(1), March 2019.

[19] Jelle Slager, Rieza Aprianto, and Jan-Willem Veening. Deep genome annotation of the opportunistic human pathogen streptococcus pneumoniae d39. Nucleic Acids Res, 46(19):9971–9989, 2018.

[20] Dominika Trzilova and Rita Tamayo. Site-specific recombination - how simple dna inversions produce complex phenotypic heterogeneity in bacterial populations. Trends in Genetics, 37(1):59–72, 2021.

