## Supplementary for "PaReBrick: PArallel REarrangements and BReakpoints identification toolkit"

### Supplementary Data for PaReBrick: Parallel Rearrangements and Breakpoints identification toolkit

#### Appendix A Parallelism Score

##### Appendix A.1 Motivation for introducing parallelism score

To explain the idea of characters' ranking based on their parallelism, we consider several examples of characters that are inconsistent with a phylogenetic tree (shown in Fig S1). In this section we interpret character states as colors for better readability.

- The character is diverged only in two strains, these strains are placed close to each other on short branches. This can be explained by inaccuracies in tree topology as well as independent rearrangements (Fig. S1a);
- The character is diverged in eight strains; seven of them form a clade, the eighth is an outgroup. This can be explained by two or even one event (Fig. S1b);
- The character is diverged in two strains, which are distant in the tree. This likely can be explained by two independent events at distant nodes (Fig. S1c);
- The character is diverged in fourteen strains, the pattern is mosaic. This can be explained only with several rearrangement events at distant nodes (Fig. S1d).

While all considered characters are inconsistent, the degree of such inconsistency varies. It depends on how many evolutionary events may explain this pattern, and how distant the respective nodes are in the tree.

##### Appendix A.2 Breaks score

To count the scores of parallel breaks we change all colors except *white* (edge exist) to *grey* (any other color). Thus, the appearance of *grey* represents the break of the considered edge. We also introduce the additional metrics:

1. *Parallel break score*: Inconsistency(*grey*), parallel rearrangement score for edge breaks;

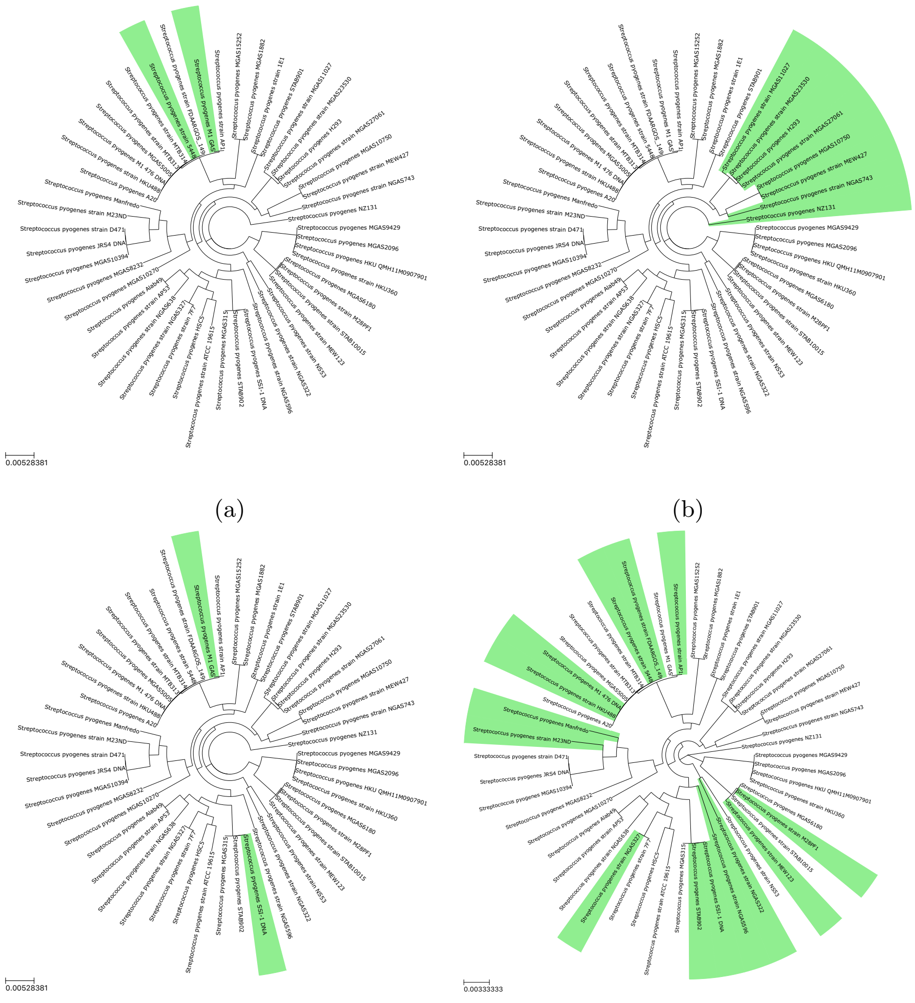

(a) (b) (c) (d)

Figure S1: Examples of parallel rearrangements for *Streptococcus pyogenes* strains

2. *Number of parallel breaks*: $|V_{grey}|$, number of independent breaks of edge.

**Example** The edge is broken in five strains out of ten in the tree (Fig. S2). According to the procedure for breaks, we switch all colors except *white* to *grey*. In this case, appearance of *grey* represents a break while *white* indicates the edge exists.

Calculation of additional metrics for estimation of breakpoint scores:

1. *Parallel break score*: Inconsistency(grey) = 4.1, parallel rearrangement score for breaks of edge;

2. *Number of parallel breaks*: $|V_{grey}| = 5$, number of independent breaks of edge.

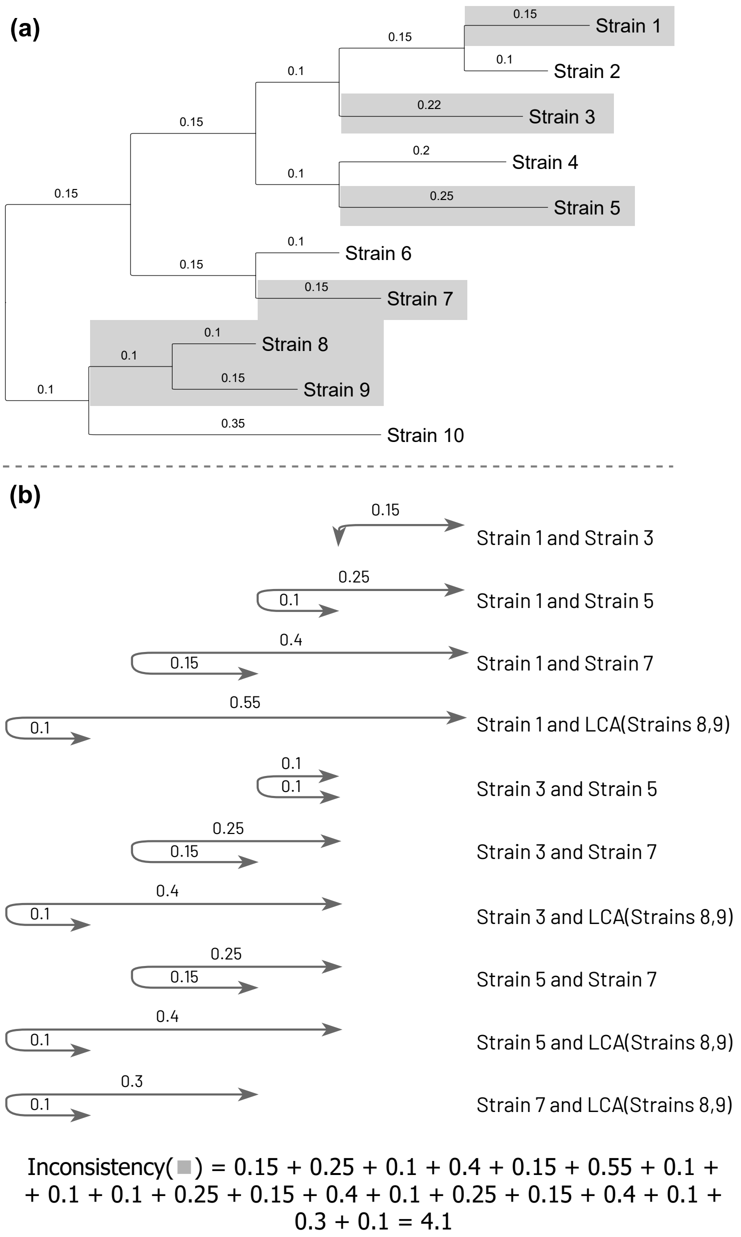


Figure S2: Same tree as in Fig. 2, all colors except *white* changed to *grey*, with calculation example for inconsistency of *grey* which represents breaks.

The PaReBrick tool uses *parallel rearrangement score* to sort the output results. As a second priority it uses *parallel breaks score* when it is applicable.

### Appendix B Example of the tool application

| block | cluster | parallel rear score | number of inconsistent colors | number of parallel events | mean copies | tree consistent |
|---|---|---|---|---|---|---|
| 166 | 0 | 6.01 | 2 | 45 | 0.38 | False |
| 159 | 1 | 4.65 | 2 | 35 | 0.35 | False |
| 157 | **2** | 4.01 | 2 | 36 | 0.43 | False |
| 156 | **2** | 3.46 | 2 | 34 | 0.36 | False |
| 155 | **2** | 3.19 | 2 | 33 | 0.36 | False |
| 172 | 3 | 2.15 | 1 | 19 | 0.11 | False |
| 2 | 4 | 1.94 | 4 | 28 | 5.92 | False |
| ... | | | | | | |

Table S1: Summary table for *unbalanced rearrangement*, only the ten first lines are shown. The blocks with the highest parallelism scores contains phage insertions; **blocks #155,156,157** are assigned to the same cluster (see 3.3). The **block #2** contains rRNA genes operon.

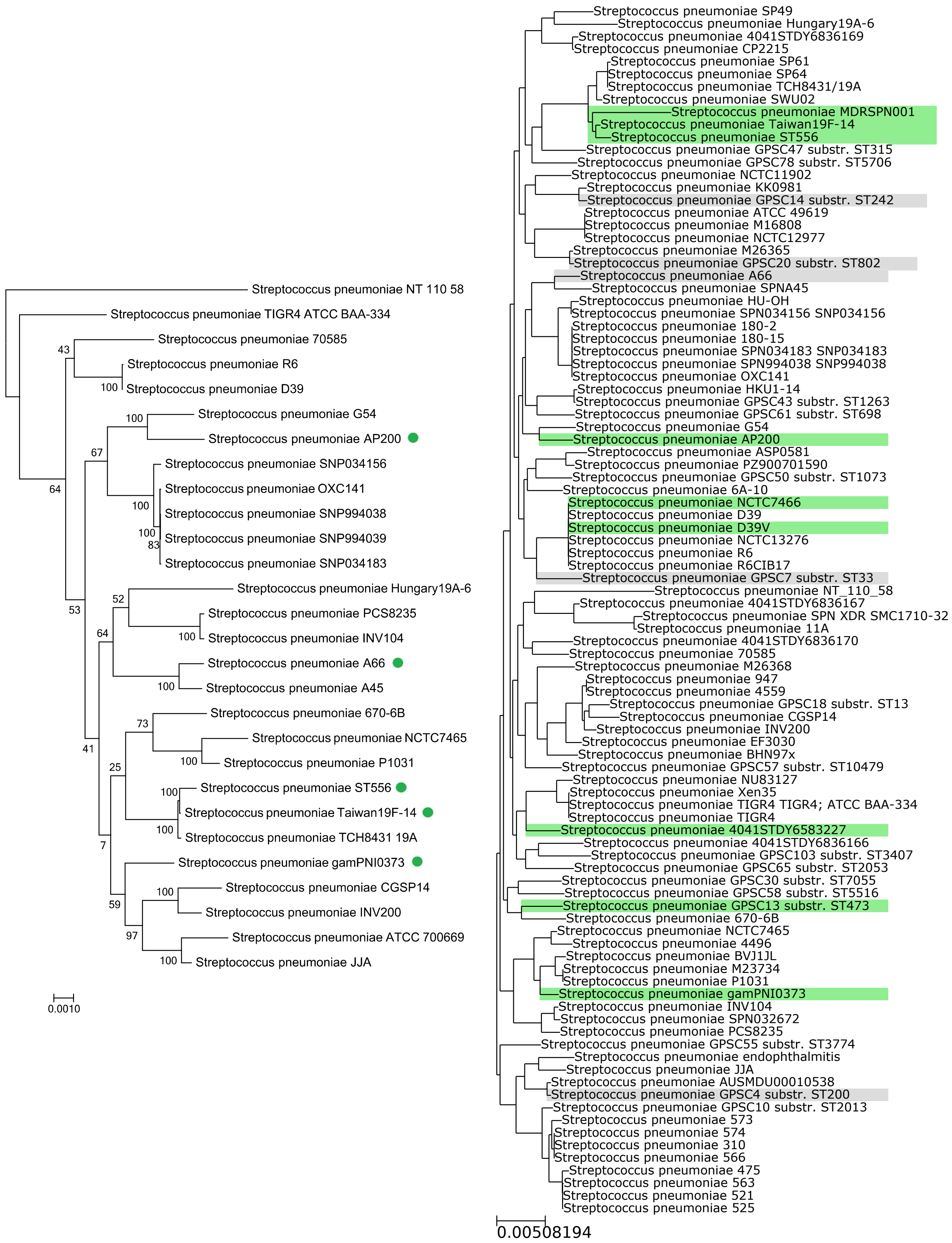


(a) the figure from the original paper. Strains with the fragment inverted are marked in green.

(b) the PaReBrick output. White represents the ancestral state of the adjacency. Strains with the fragment inverted are marked in green. Grey represents numerous rearrangements affected the adjacency.

Figure S3: Antigenic variation via large-scale inversion in *Streptococcus pneumoniae*, comparison of the PaReBrick output and the original observation.

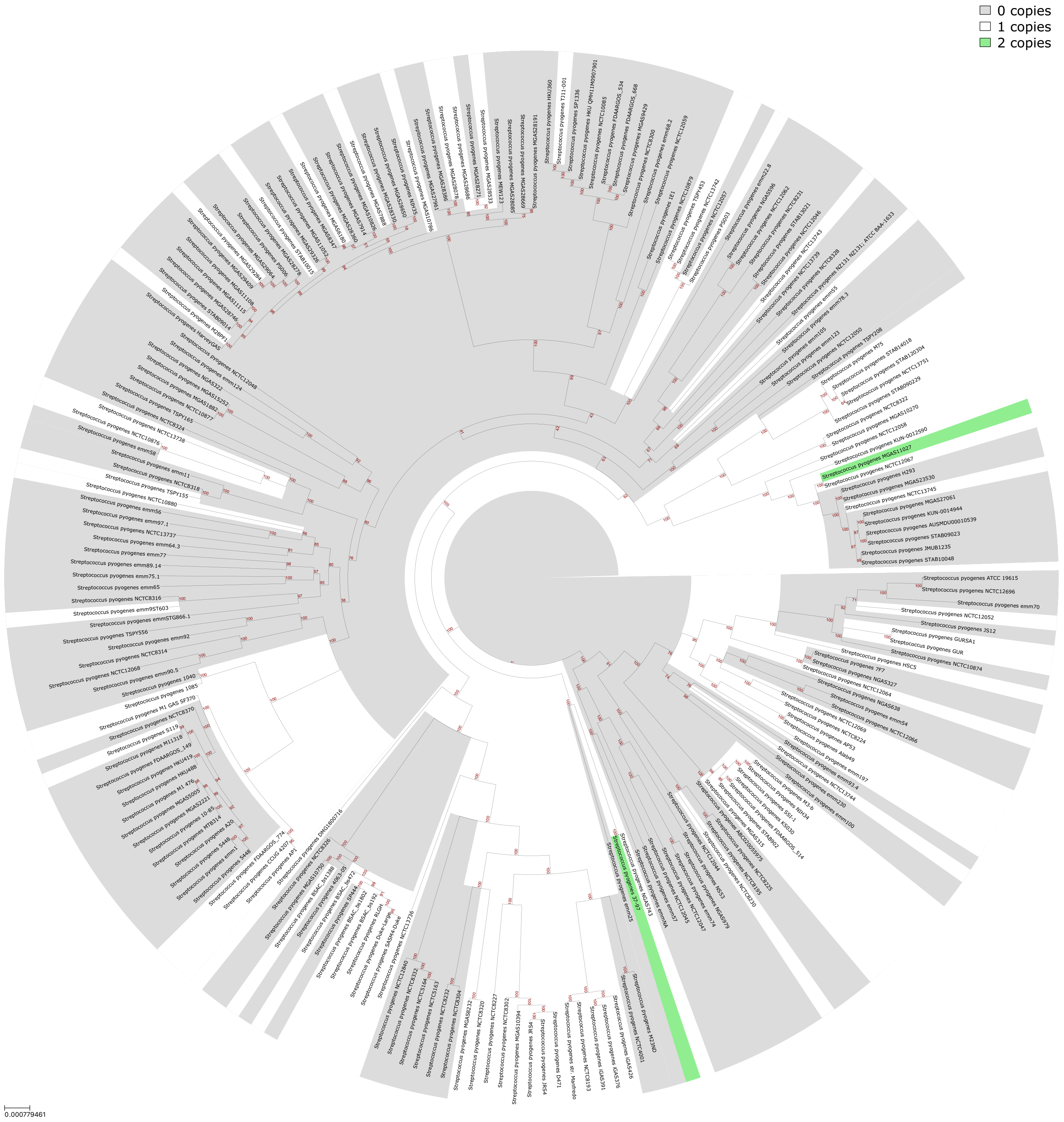


Figure S4: Copy number variation of the **block #166**. The block consists of phage genes.

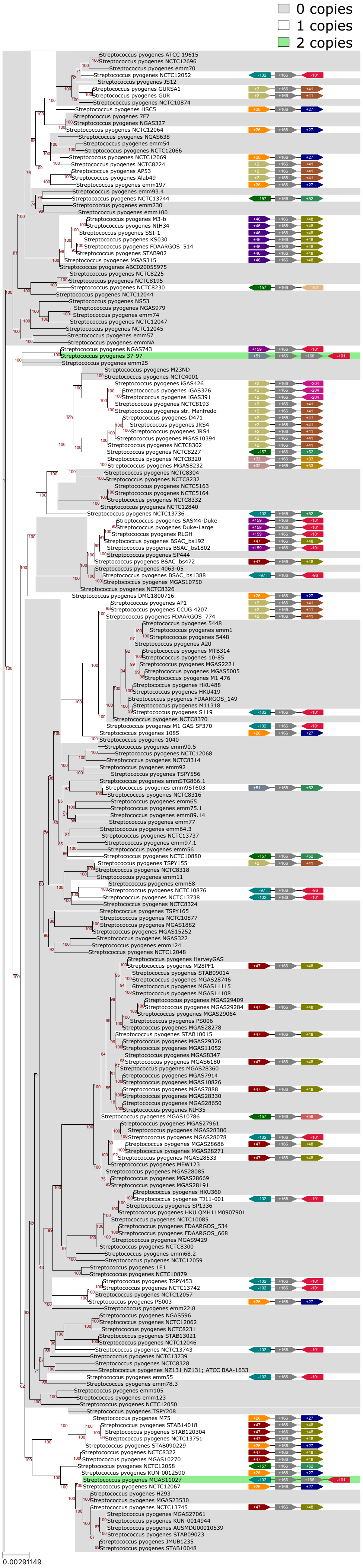


Figure S5: Genomic context of **block #166**. For each strain and each copy of the block, the upstream and the downstream neighbouring blocks are shown, revealing independent acquisition of the block by different strains.

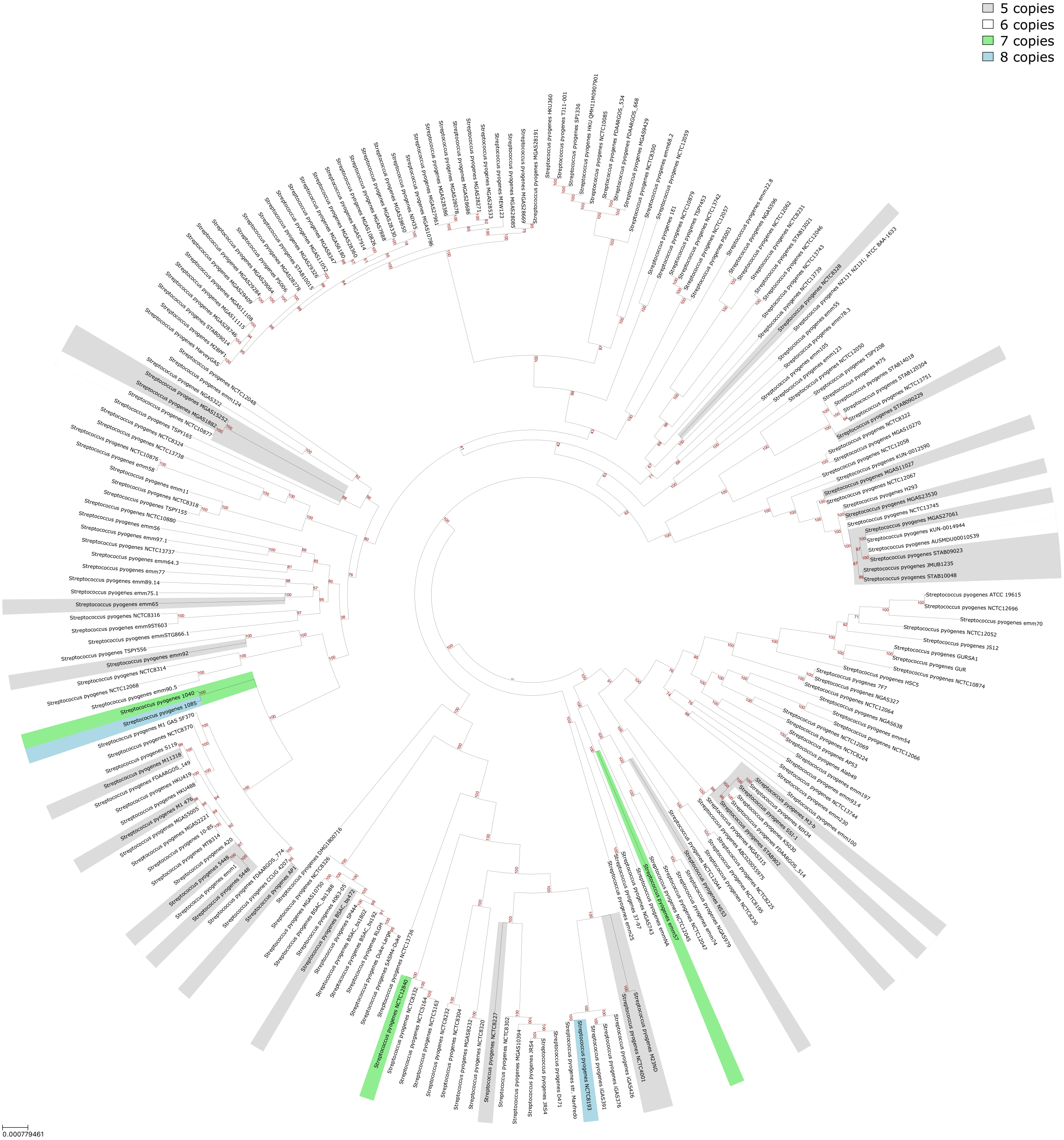


Figure S6: Copy number variation of the **block #2**. The block consists of ribosomal RNA (rRNA) genes.

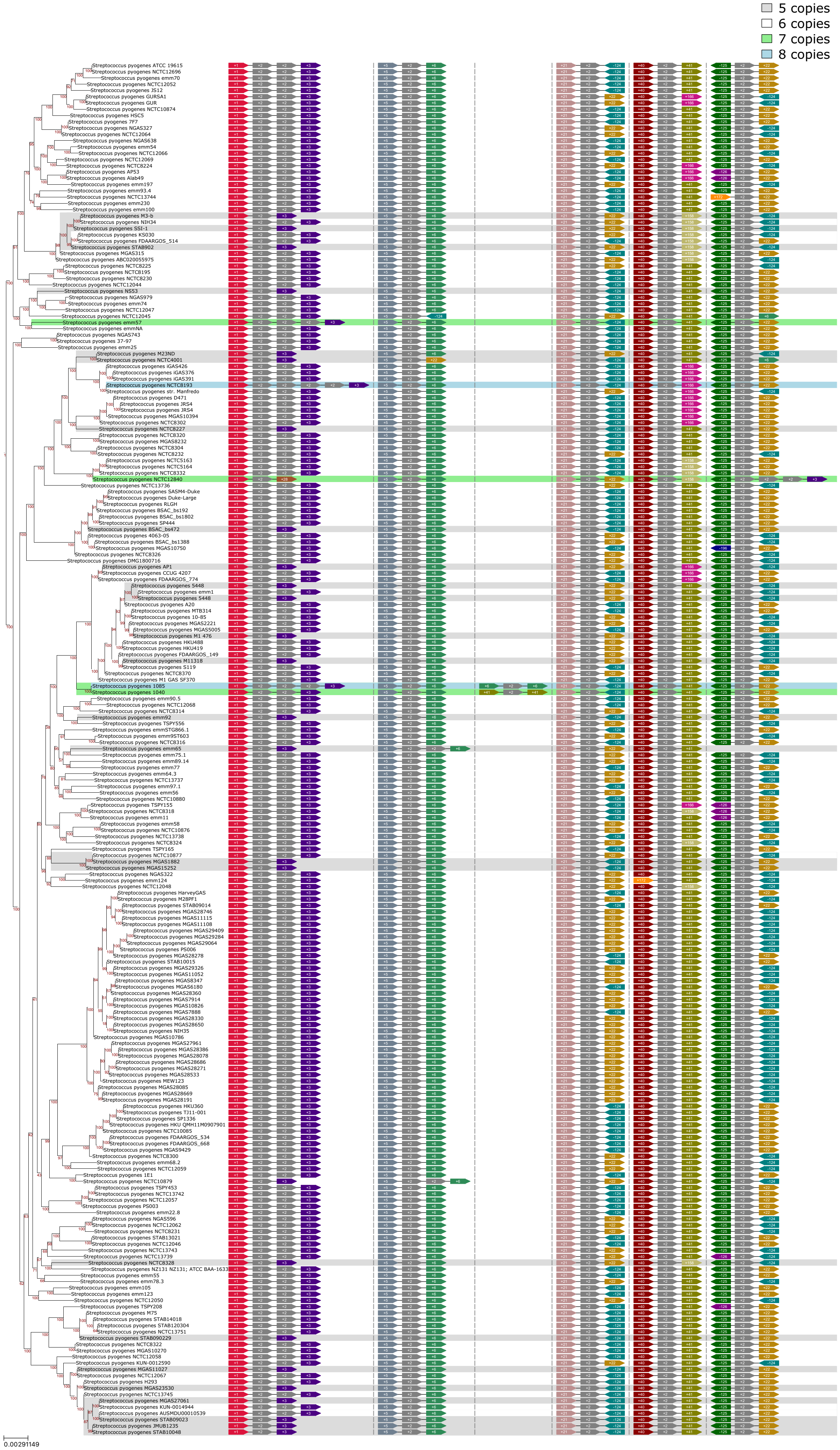


Figure S7: Genomic context of **block #2**. For each strain and each copy of the block, the upstream and the downstream neighbouring block are shown, revealing mosaic tandem duplication and numerous inversions occurred via recombination of the block's copies.

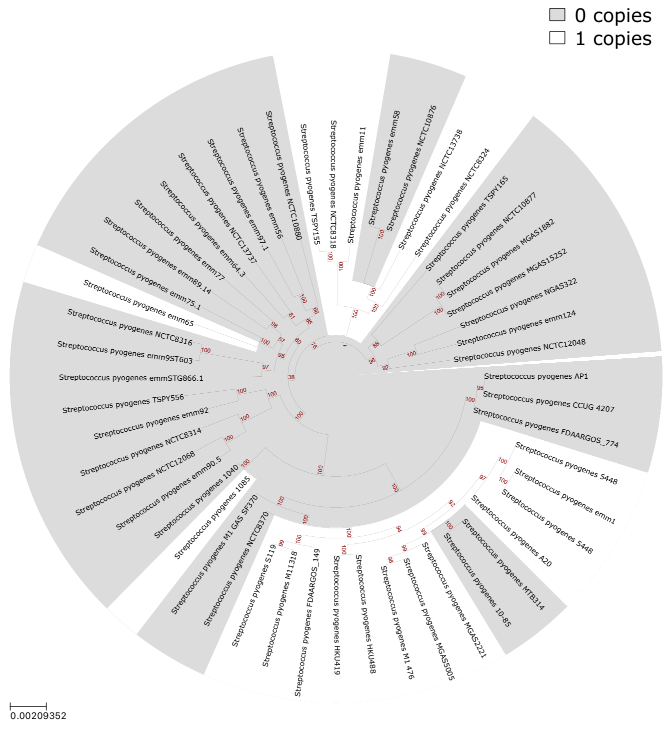


(a) Block 155

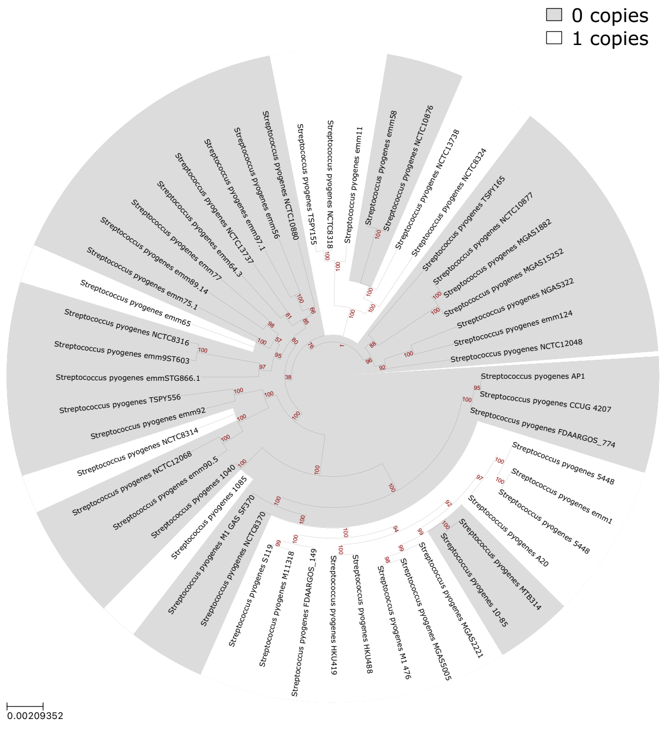


(b) Block 156

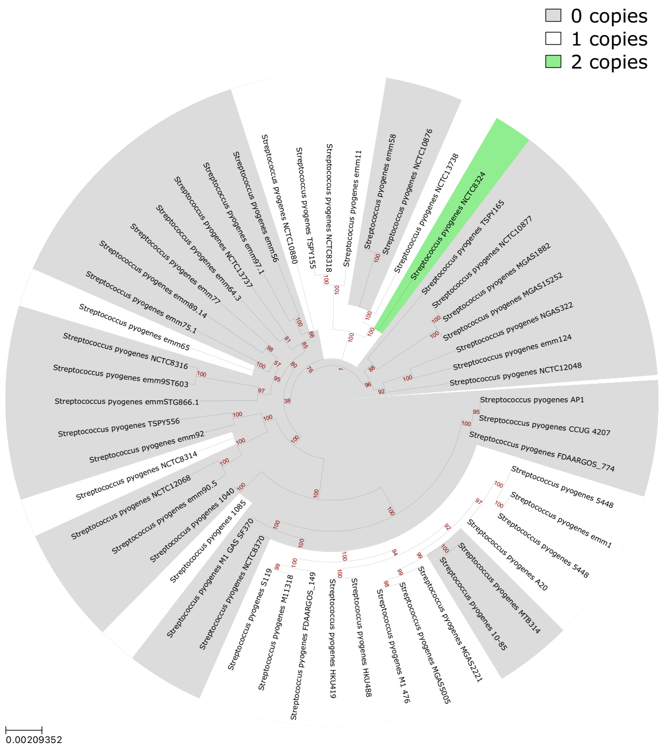


(c) Block 157

Figure S8: An example of blocks clustering, for better visibility only subtree is shown. The blocks #155, #156 and #157 are present almost in the same set of strains and placed in a row in the chromosomes. Together, the three blocks form a phage insertion.
